# Dopaminergic drug effects on probability weighting during risky decision-making

**DOI:** 10.1101/171587

**Authors:** Karita E. Ojala, Lieneke K. Janssen, Mahur M. Hashemi, Monique H. M. Timmer, Dirk E. M. Geurts, Niels P. ter Huurne, Roshan Cools, Guillaume Sescousse

## Abstract

Dopamine has been associated with risky decision-making, as well as with pathological gambling, a behavioural addiction characterized by excessive risk-taking behaviour. However, the specific mechanisms through which dopamine might act to foster risk-taking and pathological gambling remain elusive. Here we test the hypothesis that this might be achieved, in part, via modulation of subjective probability weighing during decision-making. Healthy controls (*n* = 21) and pathological gamblers (*n* = 16) played a decision-making task involving choices between sure monetary options and risky gambles both in the gain and loss domains. Each participant played the task twice, either under placebo or the dopamine D_2_/D_3_ receptor antagonist sulpiride, in a double-blind, counter-balanced, design. A prospect theory modelling approach was used to estimate subjective probability weighting and sensitivity to monetary outcomes. Consistent with prospect theory, we found that participants presented a distortion in the subjective weighting of probabilities, i.e. they overweighted low probabilities and underweighted moderate to high probabilities, both in the gain and loss domains. Compared with placebo, sulpiride attenuated this distortion in the gain domain. Across drugs, the groups did not differ in their probability weighting, although in the placebo condition, gamblers consistently underweighted losing probabilities. Overall, our results reveal that dopamine D_2_/D_3_ receptor antagonism modulates the subjective weighting of probabilities in the gain domain, in the direction of more objective, economically rational decision-making.

**Significance statement:** Dopamine has been implicated in risky decision-making and gambling addiction, but the exact mechanisms underlying this influence remain partly elusive. Here we tested the hypothesis that dopamine modulates subjective probability weighting, by examining the effect of a dopaminergic drug on risk-taking behaviour, both in healthy individuals and pathological gamblers. We found that selectively blocking dopamine D_2_/D_3_ receptors diminished the typically observed distortion of winning probabilities, characterized by an overweighting of low probabilities and underweighting of high probabilities. This made participants more linear in their subjective estimation of probabilities, and thus more rational in their decision-making behaviour. Healthy participants and pathological gamblers did not differ in their risk-taking behaviour, except in the placebo condition in which gamblers consistently underweighted losing probabilities.

## Introduction

A wealth of animal and human studies has implicated dopamine in risk-taking behaviour. Pharmacological studies in rodents have shown that drugs blocking dopamine D_1_ and D_2/3_ receptors generally decrease risk-taking, whereas drugs enhancing dopamine D_1_ and D_2/3_ receptor activity generally increase risk-taking (Zeeb et al., 2009; St Onge and Floresco, 2009; St Onge et al., 2010; Barrus and Winstanley, 2016). Similarly, in humans, boosting dopaminergic transmission with drugs such as L-Dopa and D_2/3_ receptor agonists has been shown to increase risk-taking behaviour (Riba et al., 2008; Rutledge et al., 2015; Rigoli et al., 2016; Djamshidian et al., 2010; Voon et al., 2011). Furthermore, studies in both human and animals have reported that variations in dopamine levels due to genetic manipulations or natural variations in the expression of the dopamine transporter are associated with changes in risk preferences (Mata et al., 2012; van Enkhuizen et al., 2014). Yet, the specific neurocognitive mechanisms through which increased dopaminergic transmission would increase risk-taking behaviour remain partly elusive. Some studies have suggested an influence via reward valuation mechanisms (Zhong et al., 2009) while other studies have shown that this influence is exerted via a change in value-independent gambling propensity (Rigoli et al., 2016; Rutledge et al., 2015; Timmer et al., 2017). Here we focus on a less well-investigated hypothesis, which is the role of dopamine on the subjective weighting of probabilities, both in healthy participants and individuals suffering from pathological gambling, a psychiatric disorder characterized by excessive risk-taking.

A useful and popular framework for examining how dopamine influences probability weighting is prospect theory (Kahneman and Tversky, 1979). Prospect theory posits that the departure of human agents from rational economic decision-making (i.e., expected value maximization) results from diminishing sensitivity to outcome value on the one hand, and non-linear weighting of probabilities on the other hand. People typically overweight low probabilities and underweight moderate to high probabilities, which results in an inverted-S-shaped probability weighting function and a diminished sensitivity to changes in probabilities in the medium range (Fig. 1*B*). A previous PET study in humans has shown that the degree of non-linear probability weighting in the gain domain is correlated with striatal dopamine D_1_ receptor availability across subjects (Takahashi et al., 2010). Work with fMRI has also shown that probability distortion is accompanied by similarly distorted patterns of striatal BOLD activity (Hsu et al., 2009). Here, we aimed to establish a causal link between dopamine and probability distortion using a pharmacological manipulation.

**Figure 1.**
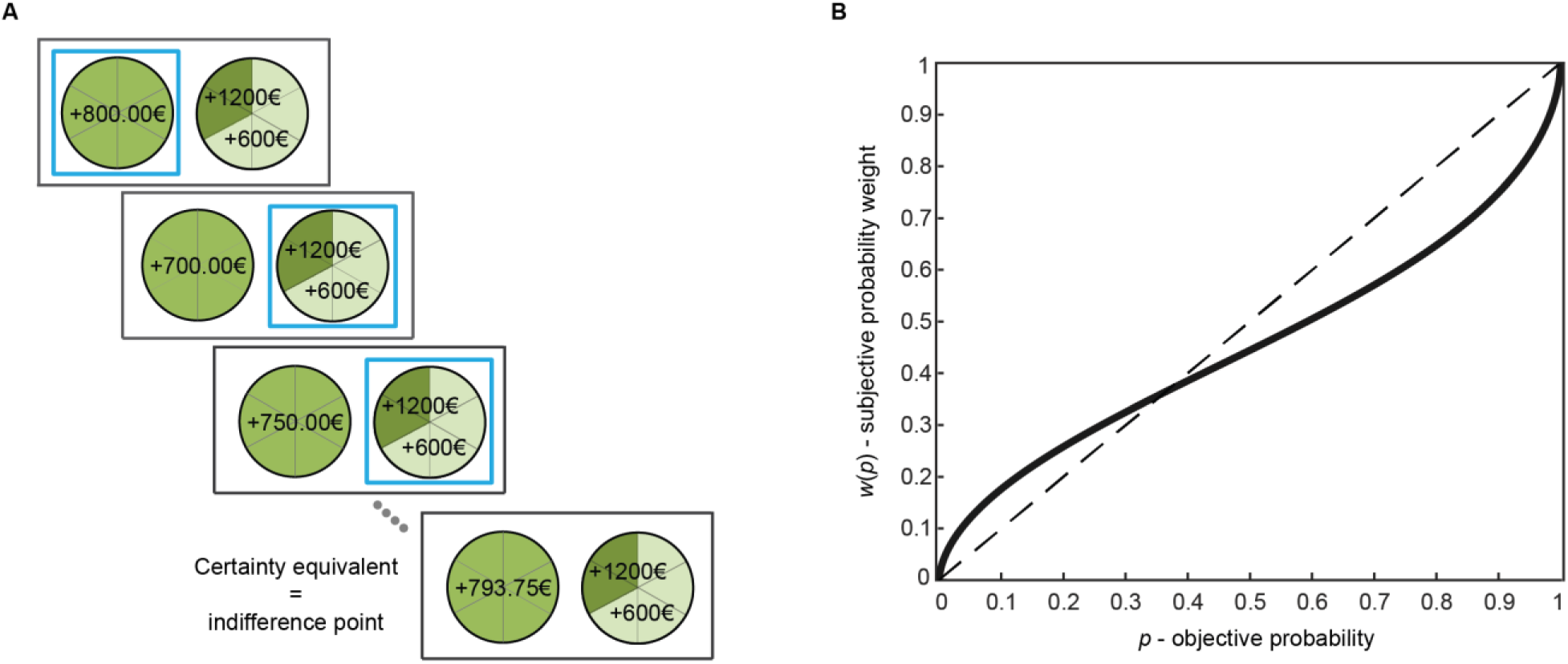
The gambling task and the probability weighting function of prospect theory. ***A***, Each trial consisted of a self-paced choice between a sure option (on the left) and a risky gamble (on the right), followed by visual confirmation of the choice (a frame around the chosen option) and fixation. The sure amount in the next trial was adjusted based on the choice (increased if gamble was chosen, decreased if the sure option was chosen), with the gamble being fixed. After six choices, the sure amount that was reached provided an indifference point between the two options, defined as the ‘certainty equivalent’ of the gamble. A new series of choices involving a new gamble was then started (in total: 10 gambles in the gain domain and 10 gambles in the loss domain). No feedback was provided on the outcome of the choices. ***B***, The solid black line represents a typical probability weighting function, with overweighting of low probabilities and underweighting of moderate to high probabilities. The dashed diagonal line represents neutrality with regard to sensitivity to probabilities.

Dopamine has been linked to pathological gambling (PG, also called gambling disorder), an addictive disorder characterized by excessive financial risk-taking in the face of negative consequences. Initial evidence came from the clinical observation that a subset of patients with Parkinson’s disease develop PG symptoms after receiving dopaminergic replacement therapy, in particular dopamine D_2_/D_3_ receptor agonists with high affinity for D_3_ receptors (Voon et al., 2009; Seeman 2015). This concurs with recent evidence showing that PG is characterized by a hyper-dopaminergic state (Boileau et al., 2014; van Holst et al., 2017), and the prominent role of dopamine D_3_ receptors in human and rat models of PG (Lobo et al., 2014; Payer et al., 2013). However, the specific mechanisms through which dopamine D_2/3_ receptor activity may act to foster PG remain elusive. In our previous study (Ligneul et al., 2013), pathological gamblers showed an elevation in their probability weighting function compared with healthy controls, reflecting an increased preference for risk or “optimism bias” in the gain domain (Gonzalez and Wu, 1999). Based on this observation, we aimed to test whether sulpiride, a selective dopamine D_2_/D_3_ receptor antagonist, could normalize risk-taking behaviour in pathological gamblers, by decreasing the elevation of subjective probability weighting.

In order to test the above hypotheses, we conducted a pharmaco-behavioural study using a within-subject, counter-balanced design. Pathological gamblers and healthy controls were asked to make choices between safe and risky options, both under placebo and sulpiride. We used prospect theory modelling to estimate subjective probability weighting and sensitivity to outcome value, separately in the gain and loss domains. Our main objective was to assess the effect of sulpiride on the two main characteristics of the probability weighting function, i.e. non-linear distortion (sensitivity to changes in probability) and elevation (optimism bias). At a more exploratory level, we were also interested in comparing those effects in the gain and loss domains, given extensive literature showing differential effects of dopamine on gains versus losses (Frank et al., 2004; Pessiglione et al., 2006).

## Materials and Methods

### Participants

We recruited 22 healthy controls and 22 pathological gamblers, all men, following an in-depth structured psychiatric interview administered by a medical doctor (MINI Plus; Sheehan et al., 1998). One gambler was excluded because his data was accidentally not written to the log file for one drug session. One control participant and five gamblers were excluded due to extreme behaviours violating core assumptions of prospect theory (see *Statistical analysis* for more details). Therefore, the reported results are based on data from 21 controls and 16 gamblers. The present task was part of a larger study for which the participants were paid €50 on each session. The other tasks in the study were a reversal learning task (Janssen et al., 2015), a slot machine task measuring sensitivity to near-misses (Sescousse et al., 2016), and a mixed gamble task measuring loss aversion. All participants provided written informed consent, which was approved by the regional research ethics committee (Commissie Mensgebonden Onderzoek, region Arnhem-Nijmegen).

Pathological gamblers were recruited through advertisement (*N* = 13) and addiction clinics (*N* = 3). None of the gamblers was in treatment at the time of testing, except for one of them who was just starting a cognitive behavioural therapy for his gambling problems. Controls were recruited through advertisement. All gamblers, with the exception of one, qualified as pathological gamblers (⩾ 5 DSM-IV criteria for pathological gambling; American Psychological Association, 2000). One gambler qualified as problem gambler as he met only four DSM-IV criteria. The severity of gambling symptoms was assessed using the South Oaks Gambling Screen (SOGS; Lesieur and Blume, 1987). All gamblers had a minimum SOGS score of 6 (range = 6–18), whereas controls, with the exception of two participants, had a SOGS score of 0 (range = 0–2).

The two groups were matched for age, net income, body mass index, and verbal IQ (Table 1). Participants were excluded if they consumed more than four alcoholic beverages daily; were using psychotropic medication; had a lifetime history of schizophrenia, bipolar disorder, attention deficit hyperactivity disorder, autism, eating disorder, anxiety disorder, or obsessive compulsive disorder; or had a past 6 months history of major depressive episode. Given the high co-morbidity between pathological gambling and other psychiatric disorders (Lorains et al., 2011), gamblers with the following co-morbidities were included: past cannabis dependence (> 5 months; *N* = 1); lifetime history of dysthymia (*N* = 1); and remitted post-traumatic stress disorder (remitted > 4 years; *N* = 1). One gambler also used cannabis weekly in the past 6 months, but did not meet the DSM-IV criteria for abuse/dependence. The control participants did not have any history of substance abuse or dependence. A number of self-report questionnaires were further used to characterize the participants (Table 1).

**Table 1.** Demographic characteristics and questionnaire scores.

| Variable | Healthy controls (*n* = 21) |  |  | Pathological gamblers (*n* = 16) |  |  |  |
| --- | --- | --- | --- | --- | --- | --- | --- |
|  | Range | *M* | *SD* | Range | *M* | *SD* | *p* |
| Age | 18–52 | 32.1 | 11.4 | 21–50 | 35.8 | 8.8 | .29 |
| Net income (€) | 0–3570 | 1691 | 1123 | 750–3250 | 1750 | 949 | .87 |
| Body mass index | 18.3–30.9 | 23.1 | 3.2 | 20.8–26.9 | 23.9 | 2.0 | .38 |
| SOGS | 0–2 | 0.2 | 0.5 | 6–18 | 12.4 | 3.9 | <.001 |
| FTND | 0–5 | 0.6 | 1.4 | 0–8 | 2.5 | 2.9 | .014 |
| Number of current smokers | – | 10 | – | – | 10 | – | .37 |
| AUDIT | 0–14 | 6.2 | 3.8 | 0–15 | 7.7 | 4.6 | .27 |
| HADS anxiety | 0–12 | 2.7 | 2.8 | 1–12 | 4.9 | 3.4 | .035 |
| HADS depression | 0–10 | 1.6 | 2.3 | 0–15 | 4.9 | 4.4 | .006 |
| Verbal IQ | 85–128 | 106 | 9.5 | 77–123 | 103 | 12.3 | .43 |

*M* = Mean. *SD* = Standard Deviation. SOGS: South Oaks Gambling Screen; FTND: Fagerstrom Test for Nicotine Dependence (Heatherton et al., 1991); AUDIT: Alcohol Use Disorders Identification Test (Saunders et al., 1993); HADS: Hospital Anxiety and Depression Scale (HADS; Zigmond and Snaith, 1983).

### Pharmacological manipulation

Participants were tested once after receiving a sulpiride pill (Dogmatil^®^, 400mg), and once after a receiving a placebo pill filled with microcrystalline cellulose. The order of administration was randomized according to a double-blind, cross-over design (placebo-sulpiride: 10 controls, 8 gamblers; sulpiride-placebo: 11 controls, 8 gamblers). The test sessions were separated by at least 1 week. Sulpiride was chosen as the dopamine-modulating drug in this study based on a few reasons. First, it is one of the most selective agents, acting selectively on dopamine D_2_/D_3_ receptors. As mentioned earlier, D_2_/D_3_ agents are known to cause pathological gambling symptoms in a subset of patients with Parkinson’s disease. Moreover, sulpiride has been shown to modulate the sensitivity to reward and punishment during learning in human studies (Eisenegger et al., 2014; van der Schaaf et al., 2014). Background neuropsychological functioning, physiological measures and subjective mood were measured at several time points during the protocol, in order to check for non-specific effects of sulpiride; no such effects were observed. The risky decision-making task was performed approximately 3 h 15 min after drug intake, thus coinciding with high plasma concentrations of sulpiride (von Bahr et al., 1991).

### Experimental design and statistical analysis

#### Experimental task

We used a “certainty equivalent” procedure (Fig. 1*A*) based on the protocol developed by Abdellaoui and colleagues (2008, 2011). Participants made series of hypothetical decisions between a sure amount of money (either a gain or a loss) and a gamble (either a pure-gain or pure-loss gamble). In each series of decisions, the gamble was fixed and the sure amount was iteratively adjusted in order to converge towards a “certainty equivalent” corresponding to the sure amount that felt subjectively equivalent to the gamble. There were 10 series of decisions (i.e., 10 different gambles) in the gain domain and 10 series of decisions in the loss domain (Table 2).

**Table 2.** Gambles with varying outcomes and probabilities.

|  | Gamble index *i* |  |  |  |  |  |  |  |  |  |
| --- | --- | --- | --- | --- | --- | --- | --- | --- | --- | --- |
|  | 1 | 2 | 3 | 4 | 5 | 6 | 7 | 8 | 9 | 10 |
| *x* | 1200 | 1200 | 600 | 1200 | 600 | 1000 | 1200 | 1200 | 1200 | 1200 |
| *p* | 1/6 | 2/6 | 2/6 | 2/6 | 2/6 | 2/6 | 2/6 | 3/6 | 4/6 | 5/6 |
| *y* | 0 | 0 | 0 | 600 | 300 | 400 | 900 | 0 | 0 | 0 |

*x* is the larger amount of money in the gamble that could be won or lost with probability *p*. *y* is the smaller amount of money in the gamble that could be won or lost with probability 1–*p*. *x* and *y* are in €. For losses, the amounts of money were the same but negative.

In each series of decisions, the sure amount offered on the first trial corresponded to the expected value of the gamble. On subsequent trials, the sure amount was adjusted based on the previous choice according to the bisection method (Abdellaoui et al., 2011), such that it was increased if the gamble was chosen, and was decreased it if the sure option was chosen. This staircase procedure drove the participants toward their “certainty equivalent”, that is, the indifference point between the risky and safe options. The decision for each trial was self-paced, after which the participant’s choice was highlighted on the screen. Participants did not receive any feedback. Each series of decisions consisted of six trials, which is considered enough to provide reliable certainty equivalent estimates (Abdellaoui et al., 2011). In order to check for errors and random responses, each series ended with two control trials that required choosing between the gamble and a sure amount slightly above or below the estimated certainty equivalent. If the participant’s response was not consistent with previous choices, the series was repeated. Participants were not explicitly informed about these control trials. We checked that the number of repetitions was not significantly different between healthy controls and pathological gamblers (gain domain: *Z* = 0.55, *p* = .60; loss domain: *Z* = 1.31, *p* = .20), between the placebo and sulpiride drug conditions (gain domain: *Z* = 1.66, *p* = .098; loss domain: *Z* = 0.36, *p* = .72), or between gains and losses in general (*Z* = 1.47, *p* = .14).

In total, participants went through a minimum of 160 experimental trials (10 series * [6 choices + 2 control trials] * 2 [gain/loss]). The task was the same in the loss domain but with negative amounts of money. Gain and loss trials were presented in separate blocks and the order of the blocks was counter-balanced across participants and drug sessions. The order of the gambles within gain and loss blocks was randomized. The task was performed on a computer and the task presentation was created with the Psychophysics Toolbox 2 (Brainard, 1997) for Matlab (www.mathworks.com).

#### Behavioural modelling

We used the semi-parametric method introduced by Abdellaoui et al. (2008, 2011; see also Fox and Poldrack, 2014) in order to estimate the value and probability weighting functions of prospect theory. This procedure was employed separately for gains and losses and for the drug and placebo conditions, within each individual participant.

In the first step of the procedure, the certainty equivalents of the gambles with varying amounts of money but a fixed probability of 2/6 (gamble indices *i* = 2,…, 7 in Table 2) were used to estimate the probability weight *w*(2/6) as well as the curvature of a parametrically defined version of the value function *v*(•). By definition, the utility of each gamble is equal to the utility of its certainty equivalent and, based on prospect theory, we can write:

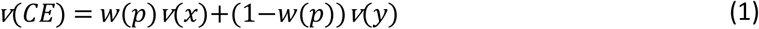

where *CE* is the certainty equivalent, *x* is the amount of money to be won with probability *p* and *y* is the amount of money to be won with probability 1-*p*. Assuming a power function *x*^*α*^ for *v*(•) (Fox and Poldrack, 2014), where *α* quantifies sensitivity to outcome values, we can further write:

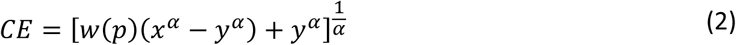

Using a non-linear least squares procedure (*lsqcurvefit* function in Matlab), we could then estimate the optimal parameter values *α* and *w*(2/6) that minimized the least squares 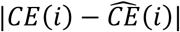, where 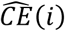 are the estimated certainty equivalents for gambles indices *i* = 2,…,7, expressed as:

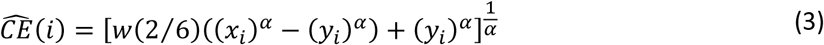

In the second step of the procedure, non-parametric estimates of the remaining probability weights *w*(1/6), *w*(3/6), *w*(4/6) and *w*(5/6) were derived from the certainty equivalents of the corresponding gambles (gamble indices *i* = 1, 8, 9 and 10 in Table 2). Since *y* = 0 in these gambles, based on equation (2) each probability weight can be calculated as follows:

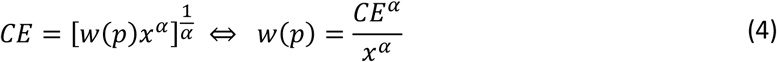

Based on those probability weights, we further derived a parametric estimation of the probability weighting function. We used a non-linear least squares procedure to estimate the two-parameter function proposed by Lattimore and colleagues (Lattimore et al., 1992), in which the sensitivity to changes in probabilities is quantified with distortion parameter *γ*, and the optimism about risk is quantified with elevation parameter *δ*:

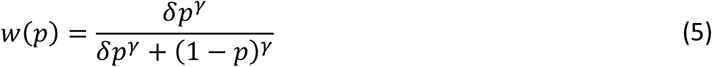

In order to avoid local minima in our least squares estimations, we used an approach with randomized starting values. The two-step estimation procedure was run 200 times with starting values randomly drawn from [0, 5] for parameters *α*, *δ*, and *γ*, and from [0, 1] for *w*(2/6). The resulting prospect theory parameters with the smallest squared norm of the residuals (‘resnorm’), reflecting the goodness-of-fit between the model and the data, were selected for the subsequent statistical analysis. Note that the ‘resnorm’ values did not differ between drugs or groups for either of the two least square estimations (paired and independent t-tests, respectively: all *p*_corr_ > 0.2), suggesting that the average goodness-of-fit was comparable across drugs and groups.

#### Statistical analysis

One control participant and four pathological gamblers were excluded from subsequent group analyses based on their certainty equivalents. Indeed, for all these participants, the absolute value of their certainty equivalent was higher for Gamble 1 (*x* = ±1200€, *p* = 1/6) than for Gamble 10 (*x* = ±1200€, *p* = 5/6), in at least one of the four conditions of interest (Gain/Loss * Placebo/Sulpiride). This behaviour violates the basic assumption of positive monotonicity in the evaluation of probabilities. One pathological gambler was further excluded due to extremely risk averse behaviour (*α* value over three standard deviations away from the mean) that likely resulted from a fear of losing control and relapsing into compulsive gambling (as reported by the participant during debriefing). While the primary analyses were performed on the reduced sample resulting from these exclusions, we also performed analyses on the full sample in order to verify that our results were not distorted by our exclusion procedure (see Sensitivity analyses in the Results section for details).

Prospect theory parameters *α*, *δ*, and *γ* were compared across groups and drug conditions, separately in the gain and loss domains, using non-parametric statistics due to the non-normal distribution of the data. Main effects of the within-subject *Drug* factor were assessed using Wilcoxon tests. Main effects of between-subject *Group* factor were assessed using Mann-Whitney *U* tests, after parameters were averaged across drug sessions. *Drug-by-Group* as well as *Drug-by-Drug Order* interactions were examined with Mann-Whitney *U* tests comparing sulpiride minus placebo values between groups. Bonferroni correction was used to correct for the six comparisons performed for each dependent variable (parameters *α*, *δ*, and *γ*): the two main effects of *Drug* and *Group* as well as their interaction, times the two contexts (gains and losses). Therefore, the corrected *p*-values correspond to the uncorrected *p*-values multiplied by 6. For effect sizes, we use the Common Language Effect sizes (CLE; Wuensch, 2015; Grissom and Kim, 2012) for intuitive interpretation. For the Mann-Whitney *U* tests, the CLE was calculated as U divided by the product of the two groups’ sample sizes. For the Wilcoxon tests, the CLE was calculated as the number of positive differences (in favour of sulpiride over placebo) divided by the number of comparisons, that is, the total sample size. Therefore, the CLE represents the probability of a randomly selected value from one group/condition being higher than a randomly sampled value from the other group/condition. For both tests, there is no difference between the groups or conditions at CLE = .5.

#### Code accessibility

The data and code used to produce the reported results are available as Extended Data. The data and code can be found with DOI references and addresses doi.org/10.6084/m9.figshare.5311354 and doi.org/10.6084/m9.figshare.5311456, respectively. The code was run with a standard Windows 7 Professional 64-bit desktop computer (Intel Xeon CPU E5-1620, 16GB RAM), both with MATLAB R2013a and R2016a.

## Results

Table 3 reports group estimates for parameters *α*, *δ*, and *γ* in the study. Figure 3 illustrates the shape of the probability weighting function separately for the gain/loss and placebo/sulpiride conditions in each group.

**Table 3.** Estimates of prospect theory parameters.

| Parameter | Controls |  |  |  | Gamblers |  |  |  |
| --- | --- | --- | --- | --- | --- | --- | --- | --- |
|  | <u>Placebo</u> |  | <u>Sulpiride</u> |  | <u>Placebo</u> |  | <u>Sulpiride</u> |  |
|  | Median | IQR | Median | IQR | Median | IQR | Median | IQR |
| $\alpha_{\text{gains}}$ | 0.74 | 0.66 | 0.80 | 0.42 | 0.80 | 0.37 | 1.16 | 0.81 |
| $\alpha_{\text{losses}}$ | 1.02 | 1.14 | 1.28 | 0.64 | 1.92 | 1.46 | 1.36 | 0.95 |
| $\delta_{\text{gains}}$ | 0.99 | 0.86 | 0.93 | 0.85 | 0.90 | 0.51 | 0.68 | 0.61 |
| $\delta_{\text{losses}}$ | 1.08 | 0.80 | 0.83 | 0.61 | 0.42 | 0.31 | 0.78 | 0.99 |
| $\gamma_{\text{gains}}$ | 0.55 | 0.50 | 0.66 | 0.54 | 0.64 | 0.35 | 0.89 | 0.49 |
| $\gamma_{\text{losses}}$ | 0.97 | 0.61 | 1.06 | 0.60 | 0.88 | 0.78 | 0.72 | 0.72 |

IQR = Interquartile range.

### Sensitivity to changes in probabilities (distortion parameter γ)

A change in the distortion parameter *γ* of the probability weighting function represents a change in the non-linear weighting and thus the sensitivity to changes in probability. The distortion parameter *γ* did not significantly differ between control participants and pathological gamblers either in the gain domain, (*Z* = 1.47, *p*_uncorr_ = .15, CLE = .64) or the loss domain (*Z* = −1.13, *p*_uncorr_ = .27, CLE = .39).

However, there was a significant effect of the drug on *γ* in the gain domain (*Z* = 2.96, *p*_uncorr_ = .003, *p*_corr_ = .018, CLE = .70). Specifically, participants had higher values of *γ* under sulpiride (*Mdn* = 0.69) than under placebo (*Mdn* = 0.58), indicating lower levels of distortion of the probability weighting function in the sulpiride condition (Fig. 2). In the loss domain, there was no difference between placebo and sulpiride (*Z* = 0.36, *p*_uncorr_ = .72, CLE = .41). Drug effect did not interact with drug order in either the gain (*Z* = 1.46, *p* = .15, CLE = .65) or the loss domain (*Z* = 0.58, *p*_uncorr_ = .58, CLE = .56), indicating no reliable session effects. The drug effect (sulpiride-placebo) was not significantly different between control participants and pathological gamblers in the gain domain (*Z* = 0.55, *p*_uncorr_ = .60, CLE = .55) or in the loss domain (*Z* = −2.02, *p*_uncorr_ = .044, *p*_corr_ = .26, CLE = .30).

**Figure 2.**
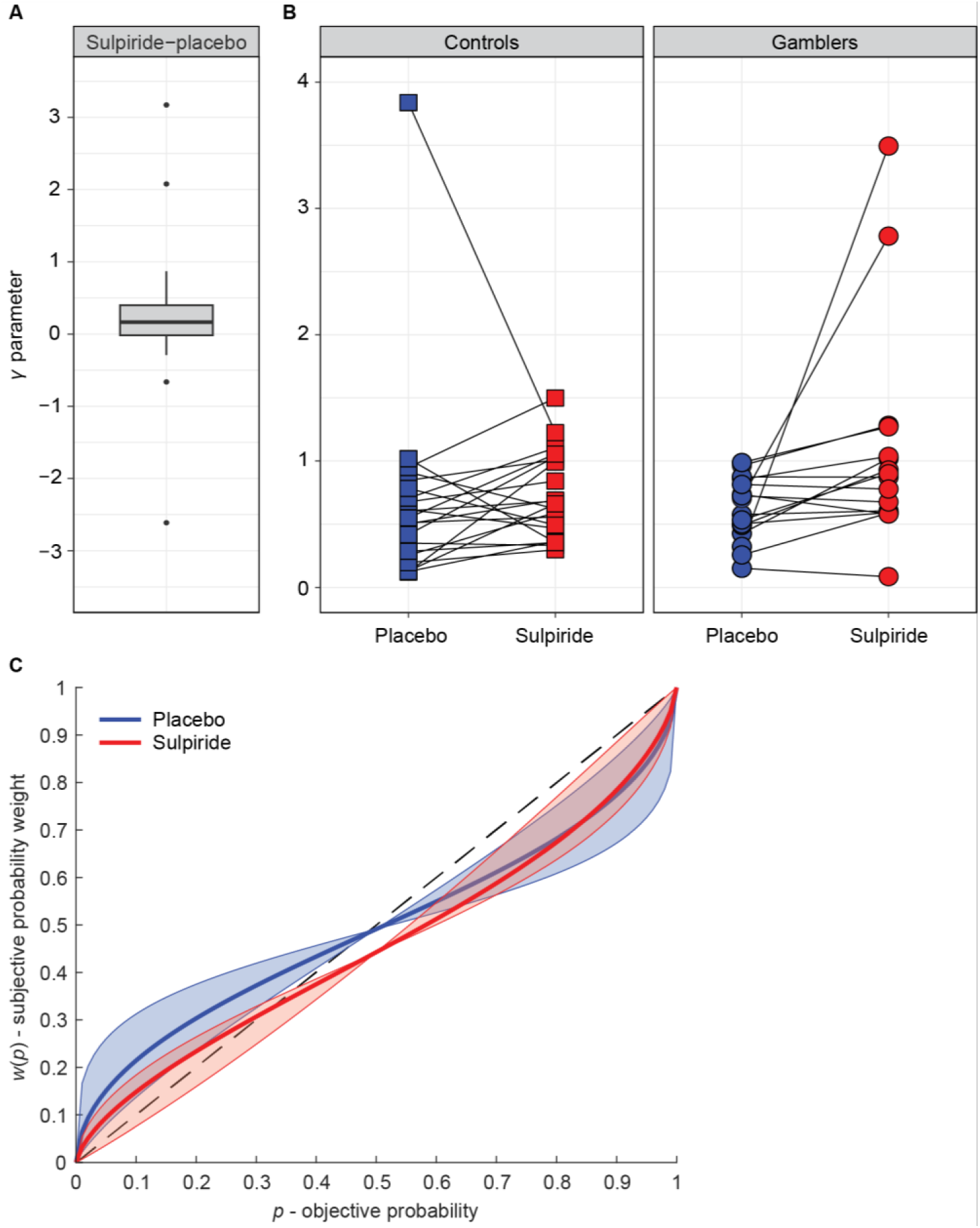
Dopaminergic modulation of probability distortion. ***A***, Boxplot illustrating the drug effect (sulpiride-placebo) on the distortion parameter *γ* of the probability weighting function in the gain domain, across all participants. Box height represents the interquartile range (IQR), black line represents the median, and whiskers represent the largest and smallest values no further than 1.5*IQR. Single data points are values are located outside the whiskers. ***B***, Within-subject paired observations of *γ* estimates in the placebo and sulpiride conditions for both experimental groups (different illustration of the result presented in Fig. 2*A*). ***C***, Fitted probability weighting function, based on the median estimates of *δ* (elevation) and *γ* (distortion) parameters across all participants. The shaded areas illustrate the variance of *γ* across participants, with the boundaries corresponding to the probability weighting function plotted with median *δ*, and 25^th^ and 75^th^ percentile *γ*.

### Optimism about risk (elevation parameter δ)

A change in the elevation parameter *δ* of the probability weighting function represents a shift in the weighting of the entire probability range, thus reflecting overall optimism or pessimism about risk. The elevation parameter *δ* did not significantly differ between control participants and pathological gamblers either in the gain domain (*Z* = −1.41, *p*_uncorr_ = .17, CLE = .36) or in the loss domain (*Z* = −1.96, *p*_uncorr_ = .051, *p*_corr_ = .31, CLE = .31). Moreover, there was no effect of drug either in the gain domain (*Z* = −0.31, *p*_uncorr_ = .76, CLE = .43) or in the loss domain (*Z* = 0.39, *p*_uncorr_ = .70, CLE = .59). Finally, the drug effect (sulpiride-placebo) was not significantly different between control participants and pathological gamblers in the gain domain (*Z* = −0.74, *p* = .48, CLE = .43) or in the loss domain (*Z* = 1.57, *p* = .12, CLE = .65).

For optimal comparison with our previous study in which we found a group difference in *δ* in the gain domain (Ligneul et al., 2013), we further compared the groups in the placebo condition alone. This analysis did not reveal a significant group difference in *δ* the gain domain (*Z* = .03, *p*_uncorr_ = 1.0, CLE = .50) but did reveal a significant difference in the loss domain (*Z* = −2.9, *p*_uncorr_ = .003, *p*_corr_ = .018, CLE = .22). Specifically, pathological gamblers had lower values of *δ* (*Mdn* = 0.42) than control participants (*Mdn* = 1.08), indicating lower elevation of the probability weighting function in the loss domain (Fig. 3*C*, *D*).

**Figure 3.**
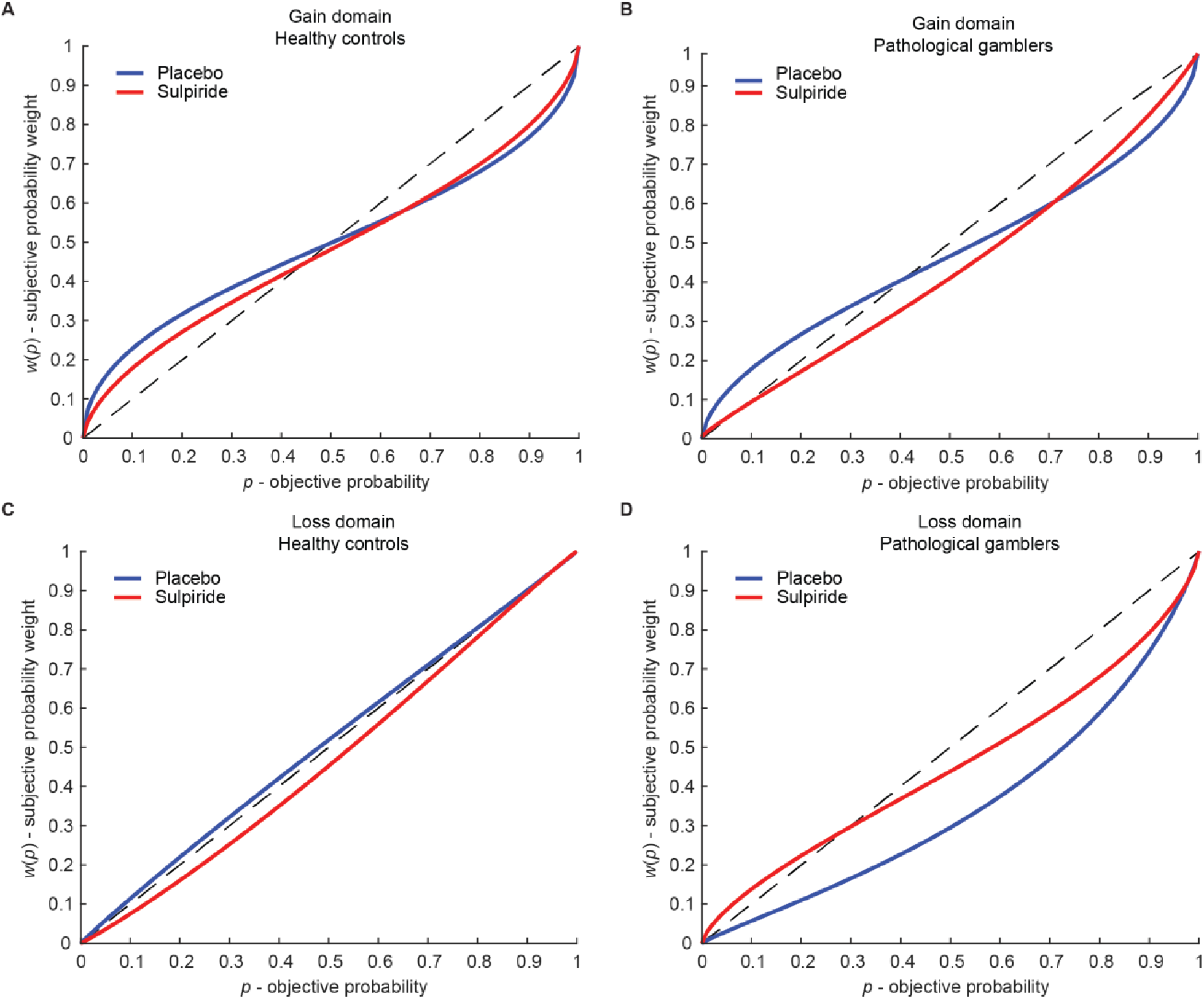
Fitted probability weighting function based on group median estimates of *δ* (elevation) and *γ* (distortion). Across groups, sulpiride decreased probability distortion in the gain domain compared with placebo (panels ***A***, ***B***). When examining the placebo condition alone, pathological gamblers showed a decreased elevation of their probability weighting function in the loss domain compared with healthy controls (panels ***C, D***).

### Sensitivity to outcomes (curvature parameter α)

Since our procedure also enabled to measure the curvature parameter of the value function, we also examined potential effects of group and drug. Non-parametric tests indicated that there was no significant difference between control participants and pathological gamblers either in the gain domain (*Z* = 0.86, *p*_uncorr_ = .40, CLE = .58) or in the loss domain (*Z* = 1.17, *p*_uncorr_ = .25, CLE = .61). Moreover, there was no effect of drug in the gain domain (*Z* = 1.53, *p*_uncorr_ = .13, CLE = .62) or in the loss domain (*Z* = −1.21, *p*_uncorr_ = .23, CLE = .41). Finally, the drug effect (sulpiride-placebo) was not significantly different between control participants and pathological gamblers in the gain domain (*Z* = 1.96, *p*_uncorr_ = .051, *p*_corr_ = .31, CLE = .69) or in the loss domain (*Z* = −1.69, *p*_uncorr_ = .10, CLE = .34).

### Sensitivity analyses

In order to confirm the pattern of our main result on probability distortion, we performed an analysis of the probability weights themselves, which were obtained using a semi-parametric procedure, as opposed to the parametric estimation of *γ*. Specifically, we performed a 2 (Groups) x 2 (Drugs) x 5 (Probability levels: 1/6, 2/6, 3/6, 4/6 and 5/6) ANOVA on the probability weights *w*(*p*) in the gain domain. We observed a significant interaction of Drug and Probability level on the *w*(*p*) (*F*(2.7, 94.495) = 3.21, *p* = .031, *η*^2^ = .084), thus strengthening our main result that sulpiride differentially modulates small versus medium-to-large probability weights. However, matched samples post-hoc t-tests between the *w*(*p*) for the two drug conditions failed to reach significance (*w*(1/6): *t*(36) = 1.15, *p* = .26, *w*(2/6): *t*(36) = 1.39, *p* = .17, *w*(3/6): *t*(36) = 0.62, *p* = .54, *w*(4/6): *t*(36) = −0.15, *p* = .26, *w*(5/6): *t*(36) = −1.41, *p* = .17).

We also repeated our estimation procedure with different variations to check the robustness of our results despite small changes in the way the parameters were estimated. First, we estimated parameters *δ* and *γ* using the Prelec version of the probability weighing function (Prelec, 1998), instead of the Lattimore version (Lattimore et al., 1992). The Prelec function is defined by the following equation:

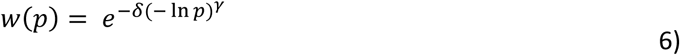

The parameters *δ* and *γ* have the same interpretation as in the Lattimore function, except that the degree of elevation decreases when the parameter *δ* increases. When the same analysis was conducted on the parameter estimates obtained with the Prelec function, the drug effect on the distortion parameter *γ* remained significant (*Z* = 2.71, *p*_uncorr_ = .007, *p*_corr_ = .032, CLE = .70), emphasizing that sulpiride decreases the distortion of the probability weighting function (i.e., increases the parameter *γ*) compared with placebo.

In addition, the drug effect on the distortion parameter *γ* remained significant (*Z* = 2.96, *p*_uncorr_ = .003, *p*_corr_ = .018, CLE = .70) when we used a linear value function (*α* = 1) instead of a power function (*x*^*α*^), a common assumption made by Ligneul et al. (2013).

Furthermore, using the original analysis with the power and Lattimore functions, the drug effect on the distortion parameter *γ* remained significant when we excluded the one participant with past cannabis dependence (*Z* = 2.83, *p*_uncorr_ = .005, *p*_corr_ = .030, CLE = .69). It also remained significant when we included all possible participants, i.e., when none of the participants were excluded based on behavioural criteria, leading to 22 healthy controls and 21 pathological gamblers (*Z* = 3.50, *p*_uncorr_ = .00046, *p*_corr_ = .0028, CLE = .72). However, the group effect on the elevation parameter previously observed in the Loss/Placebo condition did not remain significant when all participants were included, *Z* = −2.6, *p*_uncorr_ = .009, *p*_corr_ = .054, CLE = .27.

Finally, in order to assess the accuracy of the parameter estimation, we ran a parameter recovery procedure (Heathcote et al., 2015). First, we used the parameter values from the original estimation to simulate new data. Specifically, we generated synthetic certainty equivalents for every gamble (i.e., 10 gambles in the gain the domain and 10 gambles in the loss domain) for each participant and each drug condition, using equation (2). From there we created 200 noisy synthetic datasets by adding normally distributed noise to these synthetic certainty equivalents; following standards in the field, the standard error of the noise was set to be the median (over all participants and conditions) of the root-mean-squared error between the original and simulated values. We then used these noisy synthetic datasets in combination with the previously described semi-parametric procedure (Abdellaoui *et al.*, 2011), in order to perform 200 estimations of *w*(2/6), *α*, *δ*, and *γ* parameters. Across-subject correlation coefficients between the original and the recovered parameter values (defined as medians over the 200 simulations) were above .95 for all parameters in all conditions, indicating efficient parameter recovery and high accuracy in the original parameter estimation. Our main result indicating a significant drug effect on the distortion parameter *γ* showed an even larger effect size with the recovered parameter values (CLE = .76) than with the original parameters (CLE = .70).

## Discussion

This study investigated the effect of a dopaminergic manipulation on probability weighting during risk-taking in pathological gamblers and healthy participants. In line with our first hypothesis, we found that blocking dopamine D_2_/D_3_ receptors attenuated probability distortion in the gain domain. However, in contrast to our second hypothesis, the elevation of the probability weighting function was not affected by the dopaminergic manipulation and did not differ between pathological gamblers and healthy controls in the gain domain, even though a group difference was observed in the loss domain under placebo. Similarly, we did not find evidence for differences in sensitivity to outcomes between pathological gamblers and healthy controls, as well as no effect of the drug on the sensitivity to outcomes.

Our results demonstrate that the degree of non-linear probability weighting during decision-making is modulated by dopamine. More specifically, blocking D_2_/D_3_ receptors decreased probability distortion in the gain domain; this made participants more linear – or rational – in their overall assessment of probabilities, and thus more sensitive to changes in probabilities in the medium range. Such a differential effect of a dopaminergic agent on low versus high probabilities is consistent with several previous studies. For instance, Norbury et al. (2013) have shown that, in low sensation-seeking participants, the dopamine D_2_/D_3_ receptor agonist cabergoline increases risk-taking for high winning probabilities, while decreasing it for low winning probabilities. Similarly, Stopper et al. (2013) have shown that in rats, the administration of a dopamine D_1_ receptor agonist increases risk-taking behaviour in the context of high winning probabilities, but decreases it in the context of low winning probabilities. Interestingly, in all these studies including ours, the interaction of dopaminergic drug effects with probability level led to more rational behaviour maximizing long-term expected value. Thus, it could be that, instead of inducing a shift in risk-taking, modulating dopamine might induce a shift in the adherence to the principle of expected value maximization. This is an intriguing hypothesis that would deserve to be formally tested in future studies.

Particularly relevant for the current study is the work of Takahashi et al. (2010), which to our knowledge is the only study that has explicitly investigated the role of dopamine in probability weighting. In their PET study, Takahashi et al. (2010) reported that lower dopamine D_1_, but not D_2_, receptor binding in the striatum was associated with higher levels of probability distortion. This seems partly at odds with the current results, which suggest that that D_2_ receptor stimulation also plays a role in probability weighting. One hypothesis is that the drug effect observed in the current study could reflect a change in the balance between D_1_- and D_2_-receptor mediated activity in the direct and indirect pathways of the basal ganglia respectively, with sulpiride-induced D_2_/D_3_ receptor blockade being associated with a shift towards D_1_-receptor-dependent Go-pathway activity (Frank and O’Reilly, 2006; van der Schaaf et al., 2014; Jocham et al., 2011). Accordingly, we observed that D_2_/D_3_ receptor blockade decreases distortion, which is in line with the observation of Takahashi et al. that higher D_1_ receptor binding in the striatum is also associated with less distortion.

A number of previous studies have shown that dopaminergic manipulations induce a global shift in risk attitudes, i.e. they either increase or decrease risk-taking, both in humans (Rigoli et al., 2016; Rutledge et al., 2015; Djamshidian et al., 2010; Riba et al., 2008) and animals (Zeeb et al., 2009; Cocker et al., 2012; St Onge and Floresco, 2009). As mentioned previously, the lack of such an effect in our study could stem from the fact that, in contrast to most of these studies, which only manipulated one probability (or a limited range of probabilities), we considered the whole range of probabilities and observed opposite effects for high and low probabilities. Another distinctive feature of our experimental design is the absence of monetary feedback, which was meant to avoid contamination of the decision-making process by previous outcomes (Schonberg et al., 2011). This is important since risk attitudes, and in particular probability distortion, have been shown to differ when making decisions from description (as is the case in our study) versus from experience (i.e., based on feedback) (Hertwig and Erev, 2009). In addition, recent evidence in rats suggests that the influence of the dopamine D_2_ pathway on risky behaviour is exerted via the signalling of prior outcomes (Zalocusky et al., 2016). Thus, the absence of feedback in our task could explain why the blockade of dopamine D_2_ receptors failed to produce a global effect on risk attitudes. Interestingly, the vast majority of human studies reporting a global shift in risk-taking following a dopaminergic manipulation have used dopamine-enhancing agents such as L-Dopa. We are not aware of any studies reporting similar effects following dopamine D_2_/D_3_ receptor blockade.

We were not able to replicate our previous result showing an elevation of the probability weighting function in the gain domain (i.e., increased preference for risk) in pathological gamblers compared with healthy controls (Ligneul et al., 2013). One important methodological difference is that the monetary amounts used in the current study were much higher than in our previous study (300-1200€ versus 2-20€). It has been observed that people tend to be more risk-seeking for low-stake gambles than large-stake gambles, an observation referred to as the “peanuts effect” (Prelec and Loewenstein, 1991; see also Weber and Chapman, 2005). It is possible that the gamblers in our previous study were particularly sensitive to the peanuts effect and engaged in particularly high risk-seeking behaviour in the presence of low-stake gambles. It is also possible that the control participants in the current study happened to be more risk-seeking than average. The qualitative comparison of median values for the elevation parameter in the gain domain (Ligneul et al. 2013: *δ*_Controls_ = 0.74, *δ*_Gamblers_ = 1.03; current study: *δ*_Controls_ = 0.99, *δ*_Gamblers_ = 0.90) with typical values reported in the literature (Fox and Poldrack, 2014, Table A.3: median *δ* = 0.77) lends credence to these hypotheses: it seems that the control participants in the current study were more risk-seeking than average, while the gamblers were less-risking than in our previous study.

Another difference is that in our previous study, we assumed a linear value function, whereas in the current study we estimated it empirically based on the certainty equivalents. Given the trade-off between prospect theory parameters *α* (curvature parameter of the value function) and *δ* (elevation parameter of the probability weighting function) in accounting for risk attitudes (Fox and Poldrack, 2014), it could be that part of the risk-seeking behaviour was absorbed by the α parameter in our current modelling procedure, whereas all of it was absorbed by the *δ* parameter in the previous study. Note however that our present results remained qualitatively unchanged when the estimation procedure was run with a linear value function, that is, we did not observe group differences in the probability weighting function when using either the linear or power forms of the value function.

While no group difference was observed in the gain domain, analyses restricted to the placebo condition revealed that, in the loss domain, pathological gamblers showed a significant decrease in the elevation of their probability weighting function compared with heathy controls (Fig. 3*C*, *D*). This observation implies a general underweighting of losing probabilities, which could contribute to the optimism bias and excessive risk-taking behaviour observed in pathological gamblers. However, given that this result was not predicted and only applies to the placebo condition, we prefer to refrain from speculating further before it is replicated.

This study is not without limitations. First, we had a modest sample size, due partly to the complexities of running pharmacological studies in patients, and the exclusion of several participants based on outlying behaviour and violations of basic prospect theory assumptions. Yet, in order to mitigate the increased likelihood of false positives (Poldrack et al., 2017), we implemented stringent Bonferroni correction for multiple comparisons, and demonstrated the convergence of results across various sensitivity analyses. Note also that our sample was composed exclusively of men, and that further study is necessary to assess whether our results generalize to women, especially given previous evidence of gender differences in probability weighting (Fehr-Duda et al., 2006). Another limitation is the moderate test-retest reliability of decision-making measures in addictive disorders such as pathological gambling (Kräplin et al., 2016). This might have limited our ability to replicate our previous result on the elevation of probability weighting (Ligneul et al., 2013) and more generally our ability to uncover true differences between groups or drugs. Furthermore, individual risk preferences have been shown to vary substantially across tasks, a phenomenon known as the “risk elicitation puzzle” and partly attributable to inconsistent decision strategies across tasks (Pedroni et al., 2017). This observation warrants some caution regarding the generalizability of the present findings, which could be partly driven by the specific demands of the task that we used. In particular, using a more ecological gambling task might have revealed clearer differences in risk-taking between pathological gamblers and healthy controls (Schonberg et al., 2011).

In summary, this study provides evidence supporting the hypothesis that modulating dopamine affects how humans weight winning probabilities during decision-making. Dopamine D_2_/D_3_ receptor antagonism shifts probability weighting in the direction of more objective, economically rational decision-making. In future studies, it will be important to replicate this result, and further compare the contributions of D_1_ and D_2_/D_3_ receptors with the same method, since the effect has now been observed in relation to both receptors (see Takahashi et al., 2010).

## Extended Data

MATLAB code for the prospect theory parameter estimation procedure implemented in the study. The code includes scripts for the main parameter estimation, parameter recovery, and checking the quality of the estimation.

## Acknowledgements

We would like to thank Romain Ligneul, Payam Piray, Bram Zandbelt and Filip Melinščak for helpful discussions on data analysis and visualization.

